# Molecular evidence that barnacle attachment may be highly associated with a few eukaryotes in the marine biofilm

**DOI:** 10.1101/2023.10.09.561617

**Authors:** Zhizhou Zhang

## Abstract

A preliminary study on the effects of different antifouling coatings on the colonization of barnacles by investigating the diversity parameters of microflora in the coatings has been conducted. In this study, twenty-five different carbon nanotube materials were used to prepare nanocoating paints with a concentration of 0.2% w/v in iron red matrix and immersed in the sea fields for about two weeks. The number of colonized barnacles on each coating film was counted, and high-throughput sequencing analysis of their corresponding biofilm samples was performed to investigate the composition of prokaryotes and eukaryotes. It was discovered that there is no significant difference in the composition of prokaryotes in biofilms on coatings with distinct colonization levels of barnacles, but there is a significant difference in the composition of some eukaryotes such as Choreotrichia and Cryptomonadales.

## 1. Introduction

The interaction between biofilm and marine fouling organisms has been extensively studied [1a-b]; however, the different interactions vary for different foulers, resulting in no common pattern. For example, the interaction between tubeworms (polychaete Hydroides elegans) and biofilm before attachment suggests that there is a substance on the biofilm which attracts the juvenile tubeworms [1]; but barnacles seem to prefer less biofilmy surfaces; although there are also some evidences showing that certain biofilms can attract barnacles to attach [2a]. The following examples list some of the complexity of the above interactions.

Example 1: One study found that larval attachment of barnacles (B. trigonus and B. Amphitrite) was induced by biofilms developed at high temperatures (23 or 30°C), but was inhibited (B. trigonus) or unaffected (B. amphitrite) at a low temperature (16°C), suggesting a certain selectivity in the composition or surface structure of barnacles’ attachment to biofilms [5].

Example 2: Another study [6] found that barnacles prefer to attach to biofilms in intertidal zones rather than subtidal biofilms, regardless of cell density or biomass in the biofilm, indicating another selective composition and structure of the biofilm for barnacle attachment.

Example 3: This study found that some diatoms inhibited barnacles’ attachment, while others could induce barnacles’ attachment (possibly through a sugar chain structure) [8]. However, these results from pure culture-based studies may not accurately reflect the actual situation in seawater.

Example 4: The study [11] found that some compounds produced by algae Taonia atomaria can inhibit the settlement of bacteria on biofilms and inhibit barnacles larvae attachment (pure culture system).

Example 5: One study found that slides with and without seawater biofilms had different attractive forces for the tubeworm and barnacles larvae, with the latter having more attachment.

Example 6: One study found that barnacles larvae prefer to attach to biofilms from the same source (the location where barnacles larvae are collected is the same as the location of biofilm preparation), suggesting that specific adhesion signals do exist on biofilm surfaces.

Example 7: The adhesion of barnacles is complex at both molecular and community levels; the sequence structure of its adhesin belongs to a relatively special one, which is different from all other known adhesin sequences [2-4]. The relationship between barnacles’ adhesion and marine biofilm has been rarely discussed under various specific conditions, and no simple and clear conclusion has been reached.

Example 8: Complexity can be observed in another study [12], in that the promotional multispecies bacterial film turned to inhibition mode for Balanus amphitrite larval attachment in the presence of the adult extract of the barnacle, and the waterborne and the surface-associated cues from the bacteria function differentially in mediating larval metamorphosis.

Summary: There is indeed a specific relationship between barnacles’ attachment and biofilms, but it depends on specific seawater and attachment-surface conditions. It can be confirmed that barnacle larvae prefer certain cues in biofilms, but which ones need to be further identified under specific seawater and attachment-surface. An important question that has not been mentioned in previous literature is whether barnacles’ glue will attach directly to the surface of the substrate if it comes into contact with the seawater biofilm on the coating surface. A more likely scenario is that barnacles’ glue tears the contacted biofilm directly and then flows to the substrate surface beneath the biofilm, though some study did not observe apparent difference on the propensity of barnacle lava to settle on either biofilms or in their absence [13]. This aspect of research has not yet been reported; knowledge about the relationship between barnacles’ attachment and seawater biofilms remains incomplete, and further accumulation of original observations is needed to study the relationship between barnacle attachment and biofilm biology.

In this study, the authors observed whether 25 different nanomaterial-based coatings affected barnacles’ attachment to a varying degree. Results showed that at a concentration of 0.2% of nanomaterials, significant differences in barnacles’ attachment levels were observed when simply mixed with iron red anti-rust paint (Table 1). We subsequently measured the community structure of biofilm samples from these coating blocks.

**Table 1.** Basic information of nano-filled coatings, barnacle attachment and sample groupings.

| Sample | Nanomaterials No. | CNTs | Grouping<br>( prokaryotes ) | Grouping<br>( eukaryotes ) | Average attached barnacle<br>number from 3 repeats |
| --- | --- | --- | --- | --- | --- |
| WH1 | 1 | SWNT (S) | bar1 |  | ≥3 |
| WH2 | 2 | SWNT-OH (SH) | bar1 |  | ≥3 |
| WH3 | 4 | Short-SWNT (SS) | bar1 | b1 | ≥3 |
| WH4 | 18 | Short-DWNT-COOH (SDC) | bar1 | b1 | ≥3 |
| WH5 | 40 | Short-MWNT (SM5) | bar1 | b1 | $\geq 3$ |
| WH6 | 41 | Short-MWNT (SM7) | bar1 | b1 | $\geq 3$ |
| WH7 | 54 | Short-MWNT-COOH (SMC8) | bar1 | b1 | $\geq 3$ |
| WH8 | 57 | Large Inner Diameter CNTs | bar1 | b1 | $\geq 3$ |
| WH9 | 60 | Graphitized MWNT-OH (GMH2) | bar1 | b1 | $\geq 3$ |
| WH10 | 66 | Graphitized MWNT-COOH (GMC5) | bar1 | b1 | $\geq 3$ |
| WH11 | 73 | Ni coated MWNT (NM7) | | | $\geq 3$ |
| WH12 | 76 | Industrial Hydroxyl Multi-Wall Carbon Nanotubes | bar1 | b1 | $\geq 3$ |
| WH13 | 11 | High Purity Single-Walled Carbon Nanotubes | bar2 | b2 | 0 |
| WH14 | 13 | DWNT (D) | bar2 | b2 | 0 |
| WH15 | 16 | Short-DWNT (SD) | bar2 |  | 0 |
| WH16 | 17 | Short-DWNT-OH (SDH) | bar2 | b2 | 0 |
| WH17 | 29 | MWNT-OH (MH7) | bar2 | b2 | 0 |
| WH18 | 49 | Short-MWNT-COOH (SMC1) | bar2 |  | 0 |
| WH19 | 51 | Short-MWNT-COOH (SMC3) | bar2 | b2 | 0 |
| WH20 | 63 | Flash-ignited Carbon Nanotubes | bar2 | b2 | 0 |
| WH21 | 64 | Graphitized MWNT-OH (GMH8) | bar2 | b2 | 0 |
| WH22 | 65 | Graphitized MWNT-COOH (GMC3) | bar2 | b2 | 0 |
| WH23 | 68 | Graphitized MWNT-COOH (GMC8) | bar2 | b2 | 0 |
| WH24 | 70 | Short Hydroxyl Single-Wall Carbon Nanotubes | bar2 | b2 | 0 |
| WH25 | 77 | Helical Carbon Nanotubes | bar2 | b2 | 0 |

## 2. Materials and Methods

### 2.1 Coating Formulation

Carbon steel substrates (measuring 1000 mm × 1000 mm × 3 mm) were polished using abrasive paper of different grits. Afterwards, these steel panels were repeatedly washed with double distilled water, thoroughly rinsed with 70% (v/v) ethanol, and then air dried at room temperature. A commercially available chlorinated rubber-based iron oxide red paint which was commonly used to protect the steel surfaces from corrosion, was kindly supplied by the Jiamei Company (Weihai, China). CNTs (Carbon nanotubes) (Table 1) applied in the study was purchased from TimesNano company (Chengdu, China). Each type of CNTs was blended at 0.2% (w/v) with the iron oxide red paint at 1000 rpm for about half an hour at room temperature. The pretreated big steel panel were first separated into equal 100 mm × 100 mm patches by pensil lines, coated by the paint mixed with different types of CNTs patch by patch, and then cured at room temperature for three days. The 25 type of CNTs were all represented by three patches on one surface of the big steel panel, and another (back) surface of the same big steel panel was also coated with 25×3=75 patches, and the left patches in both surfaces were treated as the control.

### 2.2 Static Ocean Exposure Assays

Marine field assays were conducted at a marina on the northwest coast of Weihai, China (Lat N 37°2512, Long E 122°1740), following the Chinese national standard (GB 5370-2007), i.e., a method for testing antifouling panels in shallow ocean submergence. The upper edge of the big steel panel was immersed into seawater at a 1.5m below the lowest tide level from a static experimental wooden bridge, so the lower edge of the stell panel was seated at 2.5m below the seawater surface.

### 2.3 Sampling the Natural Biofilms plus other related data collection

A fourteen day marine in situ experiment was performed from April 3rd to 17th, 2019 under static conditions in order to obtain biofilm samples and biofouling information. All retrieved patch replicates were thoroughly rinsed with distilled water to remove the temporarily adhered debris and attached epifauna/epiphytes (but all barnacles were kept intact during biofilm sampling). About a 8 cm × 8 cm area of each patch was meticulously sampled and totally scraped with sterile brushes. Afterwards, the biofilm samples collected from the same CNTs patches were put into one sterile 15-ml tube together as a representative of all six fouling replicates, and then they were maintained at −80°C for the subsequent analysis. Biofouling and environmental information were quantitatively collected during the biofilm sampling. The big steel panel was put back into seawater for another 18 days to collect barnacle footprint sample/data after the above biofilm sampling.

### 2.4 DNA extraction, PCR amplification and high-throughput sequencing

Total genomic DNA was extracted from the representative biofilm samples using the Ezup Column Bacteria Genomic DNA Purification Kit (Cat#: B518255, Sangon, China). The assessment and quantification of DNA quality were performed using Nano-Drop ND-1000 Spectrophotometer (Nano Drop Technologies Inc., Wilmington, DE,USA). According to the DNA concentration, the DNA extracts were diluted to 1 ng/μL by adjusting to the final volume with sterile water and preserved at −80 °C. 16S ribosomal RNA gene of distinct regions (i.e. V3-V4 region) characterized from the extracted genomic DNA was amplified using the barcode-indexed bacterial primers, i.e. 338F (5’-CCT AYG GGR BGCASC AG-3’) and 806R (5’-GGA CTA CNN GGG TAT CTA AT-3’), usingn Phusion® High-Fidelity PCR Master Mix (New England Biolabs). The target PCR products were mixed with the same volume of 1 × loading buffer (contained SYBR green), and then monitored on 2.0% agarose gels. Purification of PCR products was performed using the Gene JET™ Gel Extraction Kit (Cat#: K069, Thermo Scientific, China). Sequencing library was constructed using Ion Plus Fragment Library Kit 48 rxns (Thermo Scientific, China) following the manufacturer’s recommendations. The assessment of the generated library quality was carried out on the Qubit@ 2.0 Fluorometer (Thermo Scientific, China). Eventually, the sequencing of these generated libraries was conducted on an IonS5™XL platform (Thermo Fisher, Waltham, MA, USA) at Novogene Co. Ltd, Beijing, China. Afterwards, the 400 bp/600 bp single-end reads were generated.

For 18S rRNA gene sequencing, the V4 regions of 18S rRNA genes were amplified from genomic DNA using the primer sets ssu0817F (5’-TTA GCA TGG AAT AAT RRA ATA GGA -3’) and 1196R (5’-TCT GGA CCT GGT GAG TTT CC-3’). The PCR products were then purified using a PCR purification kit (Cat#: K069, Thermo Scientific, China). The PCR libraries were constructed using Ion Plus Fragment Library Kit 48 rxns (Thermo Scientific, China). After quantification using the Qubit@ 2.0 Fluorometer (Thermo Scientific, China), the resultant PCR libraries were sequenced on the IonS5™XL platform (Thermo Fisher, Waltham, MA, USA) at Novogene Co. Ltd, Beijing, China. All sequencing data can be requested from the author.

### 2.5 Coating Characterization

#### 2.5.1 Contact Angle Measurements

The static WCA measurement was conducted on a JGW-360B Contact Angle analyzer at room temperature by depositing a water drop on each coating surface based on the sessile drop technique. In brief, when the water contacted the surface of the substrate, the syringe was moved up, leaving the droplet on the surface without motion, thus obtaining static contact angle values [23]. These tested surfaces were carefully cleaned in deionized water prior to testing. Four different points on each coating surface were tested in order to obtain an average value.

#### 2.5.2 Scanning Electron Microscopy Analysis

The surface nanostructures of selected coatings were observed on a Hitachi S-4800 Scanning Electron Microscope (SEM, Hitachi Limited, Japan), which was operated at the acceleration voltage of 15.0 kV. All of the coating samples were sputter-coated with gold prior to characterization in order to minimize sample charging.

### 2.6 Data Analysis

Sequencing analysis was conducted in the Quantitative Insights into Microbial Ecology framework (QIIME, Version 1.9.1) as Kuczynski et al described earlier [17]. Briefly, the Cutadapt (Version 1.9.1) [18] software was applied to initially screen and remove the low-quality parts of the reads, and then the sample data was separated from the obtained reads according to their unique barcodes. The raw reads were obtained by removing the barcode and primer sequence. The identification of chimeric sequences was conducted to compare the reads with Silva database using UCHIME algorithm [19-20]. Then the final clean reads were acquired by cutting off the chimeric sequences therein. All clean Reads for all samples were clustered using UPARSE software (version 7.0.1001) [21], which were clustered into Operational Taxonomic Units (OTUs) with a 97% identity. The sequence with the highest frequency of occurrence in the OTUs was screened as a representative OTU sequence for further annotation according to the algorithm. Taxonomic annotation of representative OTUs was performed based on Silva’s SSUrRNA database (set threshold of 0.8∼1) via the Mothur algorithm [22]. OTUs abundance information was normalized using a standard of sequence number corresponding to the sample with the least sequences. The subsequent alpha and beta diversity analysis were implemented according to this output normalized data.

## 3. Results and Discussion

For 25 biofilms, 25 were successfully for 16S rDNA amplification and further sequencing, but only 20 were successfully for 18S rDNA amplification and sequencing. The 25 16S samples were grouped into bar1 and bar2 according to their barnacle attachment average numbers, while the 20 18S samples were grouped into b1 and b2 accordingly (Table 1).

The barnacle-rich group (bar1) and barnacle-poor group (bar2) had very similar profiles of bacterial community (Figure1A), only with apparent ratio difference between two genera: Cycloclasticus and norank_o_Chloroplast (without the genus name). Cycloclasticus has a function of degrading polutants in the ocean [16] and is a marker as ocean pollution. For eukaryotes community, b1 and b2 had much larger difference than that between bar1 and bar2, especially the main differences existed in the top 10 dominant genera (Figure1B). Among the top 10 eukaryotic genera, actually only Leucocryptos, Salpingceca, Nuclearia, and Bradymyces have genus names. This result is consistent with the previous finding that eukaryotic has a larger difference than prokaryotic communities on the surfaces of different nano-filled coatings [14a-b]. The reasons why eukaryotes are more sensitive to nano-filled coatings than prokaryotes are still vague. The potential explanation is that eukaryotes normally are larger in cell size, and thus easier to be interfered by some nano-spikes on a surface. This kind of interference would finally affect spatial pattern and composition of eukaryotes on the coating surface. This guess may be confirmed, in the future, by prokaryotes with big difference in cell sizes. But whether and how the big difference on eukaryote compositions between b1 and b2 directly contributes to barnacle settlement descrepancy is still a question. Importantly, such a big composition difference between b1 and b2 was not demonstrated by diversity indices (supplementary data not shown). Results in Figure 1 were well represented in Figure 2, in which only eukaryotes showed apparent taxonomic differences with significance in Choreotrichia proportionally correlated with the adhesion degree of barnacles while Cryptomonadales negatively correlated.

**figure 1.**
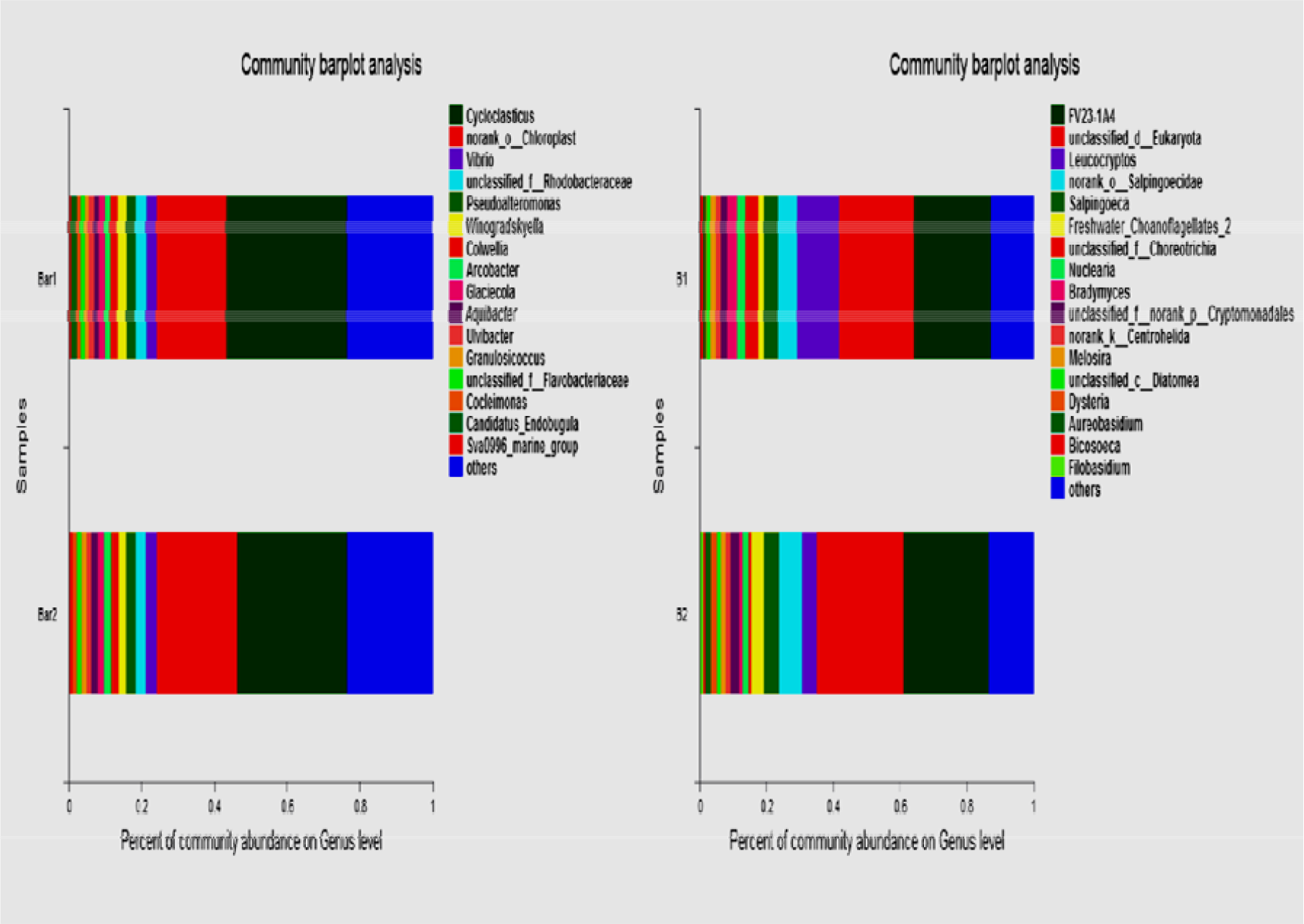
prokaryotic (a) and eukaryotic (b) community compositions compared between barnacle-rich and barnacle-poor groups.

**Figure 2.**
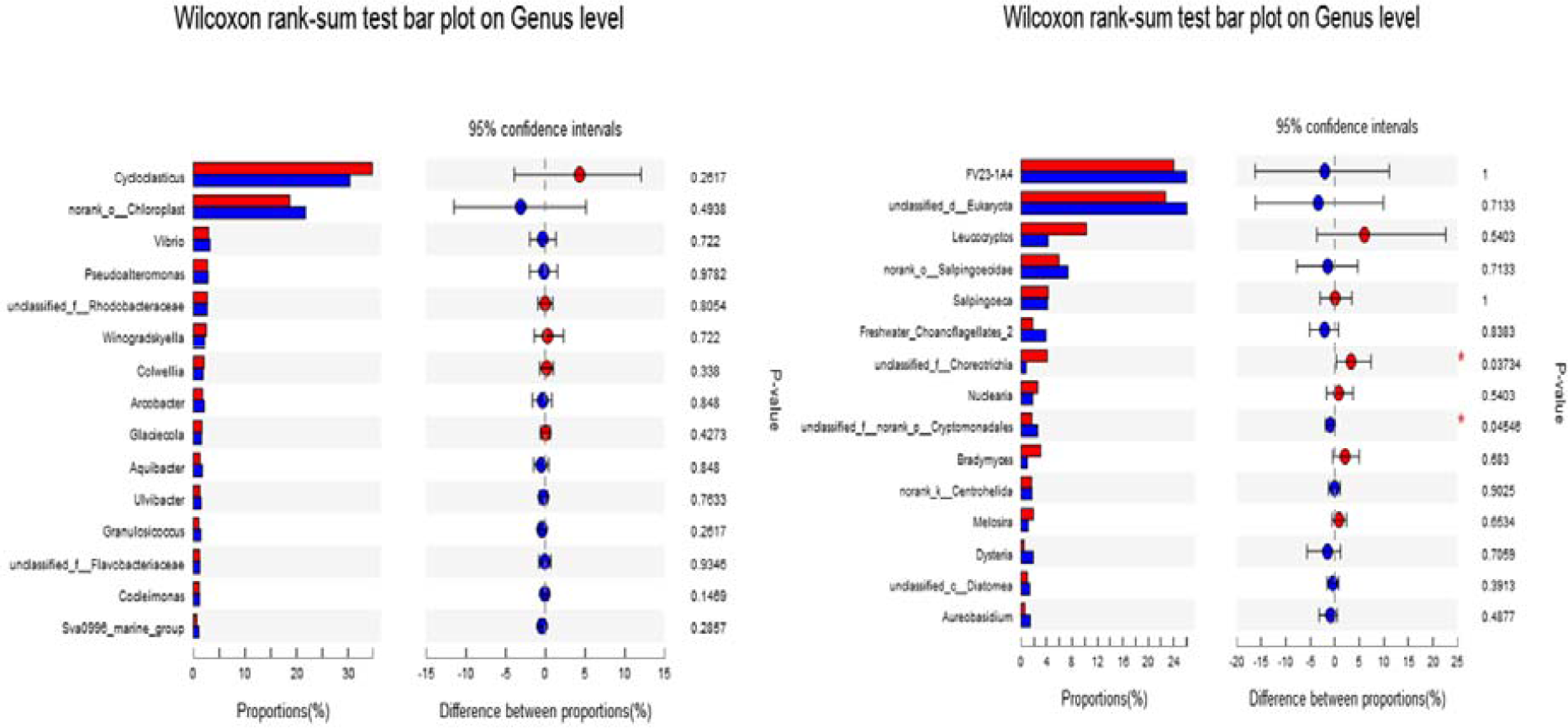
The microbial compositional structures between barnacle-rich coatings and barnacle-poor coatings are highly similar in prokaryotic, but significantly different in eukaryotic (for several taxonomic units) communities. Red color represents bar1 or b1, and blue color represents bar2 or b2 in Table 1.

The RDA analysis (Figure 3) used basic parameters of nanomaterials and antifouling performance parameters of each coating (the number of each type of macrofouler in a patch), including barnacles (Bar), black tubeworms (Black), sea squirt (ascidians, HQ), grass snails (bryozoans,Tai) and total antifouling score (AF); the parameters of nanomaterials included outer diameter (OD), dispersion (DP), length (L) and specific surface area (SSA). The prokaryotic RDA analysis showed that the layer was apparently divided into two categories, one positively correlated with chloroplast content and the other positively correlated with Cycloclasticus; however, the antifouling score AF seemed to be not closely related to these two genera; while the eukaryotic community showed a significant difference in the correlation between nanomaterial parameters. For example, SSA had a relatively small correlation with the prokaryotic community (the arrow of SSA was very short), but had a strong negative correlation with the number of barnacles; however, on the eukaryotic side, SSA showed a strong positive correlation with most sample communities, and had almost no correlation with the number of barnacles; In addition, the number of barnacles showed a very small correlation with the antifouling score in the prokaryotic map, but there was a strong positive correlation between the number of barnacles and the antifouling score in the eukaryotic map (this correlation can be quantitatively characterized). There were also huge differences between the results of prokaryotic and eukaryotic RDA analyses in terms of correlations between nanomaterial parameters themselves, such as the correlation between Tai and L, which was almost zero in prokaryotic and showed a huge negative correlation in eukaryotic. Moreover, Tai showed a huge negative correlation with several nanometric parameters on the eukaryotic map.

**Figure 3.**
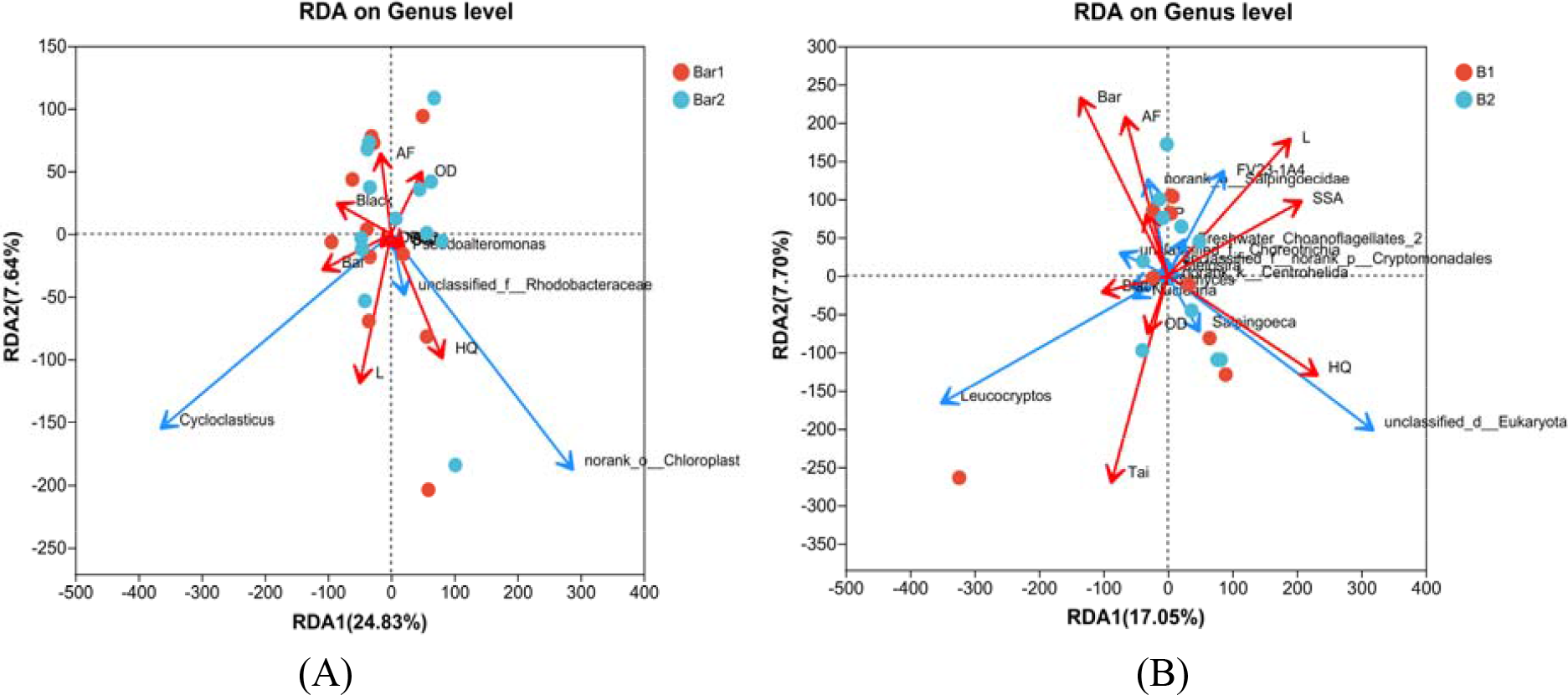
RDA analysis of microbial community structures and coating parameters in Table 2.

**Table 2.** Coating and environmental parameters used for correlation analysis.

| Sample | OD | L | DP | SSA | HQ | Black | Tai | Bar | AF |
| --- | --- | --- | --- | --- | --- | --- | --- | --- | --- |
| WH1 | 1.5 | 17.5 | 3 | 380 | 5.2 | 6 | 2.1 | 2.5 | 79.1 |
| WH10 | 25 | 20 | 4 | 55 | 9.2 | 4 | 3.9 | 1 | 76.8 |
| WH12 | 20 | 30 | 3 | 100 | 9.2 | 4 | 3.9 | 3 | 87.3 |
| WH13 | 2 | 17 | 3 | 380 | 3.7 | 4.8 | 10.2 | 2 | 74.3 |
| WH14 | 3 | 25 | 5 | 350 | 8.4 | 6.8 | 3.7 | 1.3 | 74.9 |
| WH15 | 3 | 1.3 | 4 | 350 | 7.2 | 6.9 | 4.2 | 1 | 75.7 |
| WH16 | 3 | 1.3 | 4 | 350 | 4.8 | 6.5 | 4.4 | 1.5 | 77.9 |
| WH17 | 40 | 10 | 4 | 60 | 7.2 | 3.5 | 6 | 0.5 | 77.8 |
| WH18 | 4 | 1.3 | 3 | 500 | 5.8 | 11.3 | 4.2 | 0.3 | 73.5 |
| WH19 | 15 | 1.3 | 4 | 200 | 10.6 | 4.8 | 7.2 | 0.5 | 72 |
| WH2 | 1.5 | 17.5 | 5 | 380 | 8.8 | 4 | 3 | 2.5 | 76.6 |
| WH20 | 25 | 25 | 5 | 300 | 0.4 | 7 | 2.4 | 2.5 | 82.7 |
| WH22 | 15 | 15 | 4 | 100 | 2 | 18.8 | 2.2 | 1 | 71 |
| WH21 | 50 | 15 | 4 | 20 | 2.5 | 4.3 | 2.4 | 1.8 | 84.1 |
| WH23 | 50 | 15 | 4 | 20 | 3.3 | 13.2 | 3.8 | 0.8 | 73.9 |
| WH24 | 1.5 | 2 | 4 | 380 | 6.8 | 2 | 3.8 | 1.8 | 80.7 |
| WH25 | 150 | 5.5 | 1 | 30 | 6.8 | 2 | 3.8 | 1 | 86 |
| WH26 | 1.5 | 17.5 | 5 | 380 | 8.9 | 4 | 3 | 2.5 | 76.6 |
| WH3 | 1.5 | 2 | 5 | 380 | 6.1 | 4.3 | 9.4 | 2.8 | 72.5 |
| WH4 | 3 | 1.3 | 4 | 350 | 10.9 | 1.4 | 6.2 | 2 | 74.5 |
| WH5 | 25 | 1.3 | 4 | 110 | 4.6 | 5.8 | 6.8 | 2 | 75.9 |
| WH6 | 40 | 1.3 | 4 | 60 | 8.2 | 9.3 | 4.4 | 2.3 | 70.9 |
| WH7 | 50 | 1.3 | 4 | 40 | 3.3 | 3.8 | 3.7 | 1.8 | 82.5 |
| WH8 | 45 | 35 | 5 | 200 | 3.9 | 5 | 3.6 | 0.8 | 81.8 |
| WH9 | 11.5 | 25 | 3 | 117 | 15.4 | 3.3 | 4.2 | 1.5 | 70.7 |

Note: OD: outer diameter (nm); L: length(nm); DP: dispersion level (from 1 to 5); SSA (specific surface area, m<sup>2</sup>/g); HQ: average numbers of attached ascidians ; black: average numbers of attached black tubeworms; Tai: average numbers of attached bryozoans ; Bar: average numbers of attached barnacles; AF: antifouling score;

The parameter Black has positive correlation with about half samples, but in Figure3B the Black parameter has positive correlation with only three samples; The Bar parameter of most concern in this study, has similar situation as Black. In general, the RDA analysis employed seven parameters and all of them have apparent different correlation outputs between prokaryotic and eukaryotic communities in the biofilm samples.

The researches of the authors’ lab in recent years have found that under 0.1-0.2% doping concentration, CNTs nanomaterial series generally fail to bring about significant changes in the contact angle of seawater on the coating surface [14a, 23], that is, nanomaterial fails to significantly change the hydrophilicity and hydrophobicity of the coating surface. Besides, SEM results for nano-filled coatings did not display apparent morphological changes on the coating surfaces (data not shown), though there are some small spikes out of the coating surface at some local positions. However, the above results did not exclude the possibility that some CNTs may significantly change the adhesion film formation process of eukaryotic organisms at local positions, then reshape the relative distribution of different bacterial genera in the whole biofilm, which will directly affect the degree of attachment of barnacles larvae. Prokaryotic microorganisms are much smaller in size in average than eukaryotic microorganisms, and that is an important reason why any surface has prokaryotes practically capable of attachment, so they are less affected by nanomaterials on the coating surface than eukaryotic microorganisms. This is consistent with the repeated results of laboratory measurements of biofilm flora.

It is possible that the biofilm surface colonization of barnacles is highly determined by some eukaryotic microorganisms, and this speculation already has some evidence. The main findings of this study (Figure 1-2) revealed two groups of eukaryotes with significant difference in the two coatings; and the microbial composition beneath the barnacles’ footprint (attachment plaque) was also predominantly eukaryotes [15] (Table 3). In theory, a few bacterial species differences within the biofilm could have been enough to cause a large change in the distribution pattern of bacteria on the biofilm surface, which in turn would lead to whether or not a specific biofilm surface is suitable for barnacle attachment. However, research into this area has been very limited so far. If only a few bacterial species were observed in the two biofilms when using the coatings based on barnacles’ attachment levels, these bacterial species should be considered as the direct cause of the difference in barnacle attachment levels, at least closely related ones. There is also a possibility that there was time lag between barnacles attaching to different patches of the biofilms. Once an attachment event occurred, it might have caused local habitat changes and subsequently changed the degree of attachment of certain eukaryotes, resulting in differences in eukaryote composition. This is also possible. If so, it seems unlikely that most of the microbes underneath a barnacles’ imprint are eukaryotes; our current preliminary results suggest that most of them are eukaryotes. Thus, it is more likely that these eukaryotes first influenced barnacles’ attachment, or rather attracted their attachment, though it is intriguing why some Choreotrichia species were not shown in Table 3.

**Table 3.**
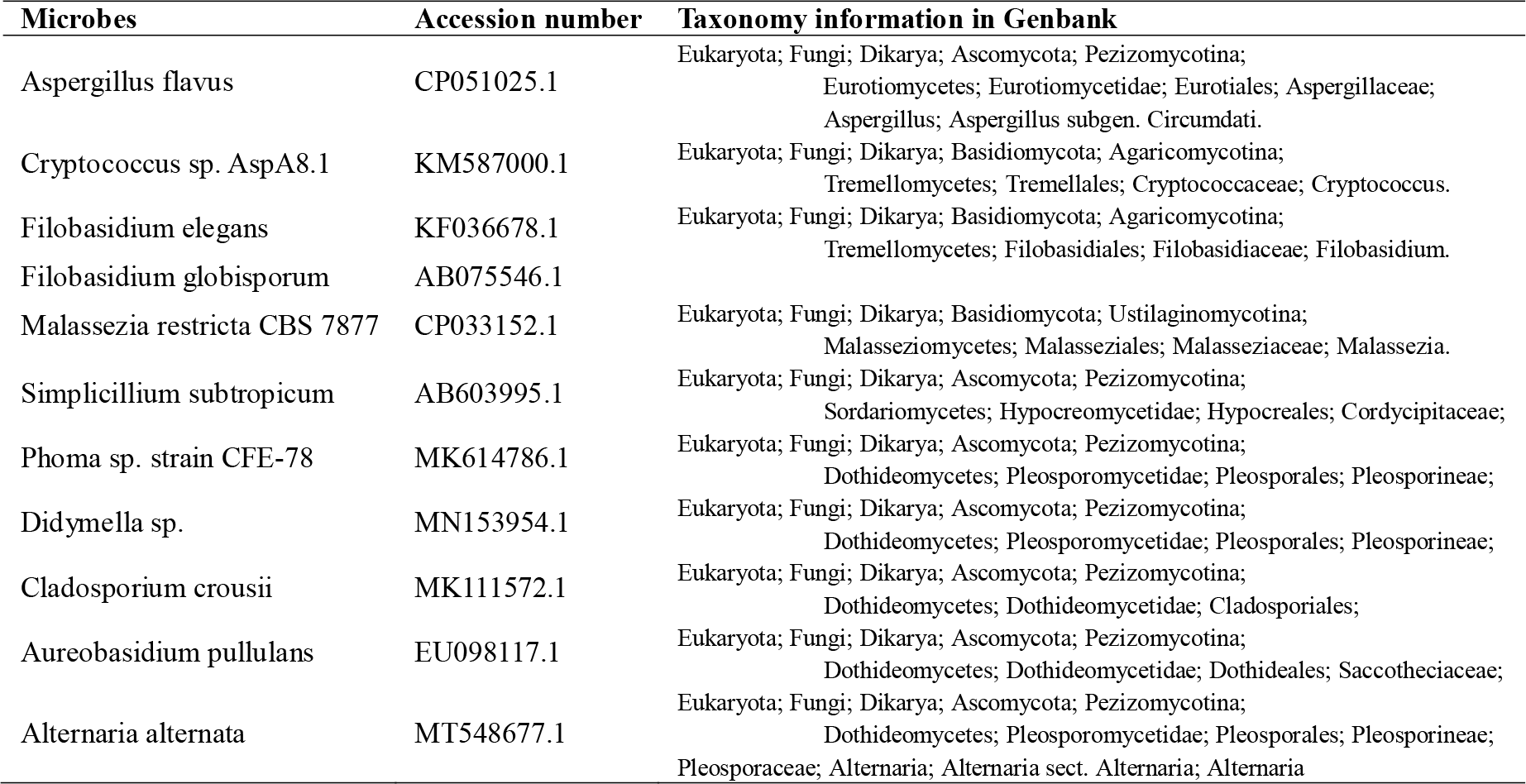
Microbes determined underneath the barnacle plaques^[15]^.

Literatures about barnacles attachment can be read to find that preference of barnacles larvae for attaching substrate is existed. This preference may be the result of billion-year evolution in order to improve the survival efficiency; or it may be that barnacles attach basically randomly, but different fates will be followed after they are attached at different locations, and some bacterial genera can increase their survival rate (due to easy access of nutrient or avoiding more predators etc.) Initially, this research found that certain eukaryote genera on biofilm might be the key point of barnacles attachment. Even though barnacle larvae’s attachment organs maybe haven’t evolved specific adsorption mechanism towards these eukaryote genera, barnacles that attach onto these eukaryote genera randomly may have more chances to survive. Although this research is not enough to get very definite conclusion, the data available remind researchers that they have to pay attention to the above possibilities. If barnacles larvae attach process to these eukaryote genera is not random, then in real seawater environment, we suppose that these eukaryote genera should release chemical attractant (as chemical inducer or even attractive food for barnacles larvae) or possess special surface adsorption structure; when barnacles larvae sucker disc contacts with such surface structure of eukaryote genera, a strong vacuum adsorption will be formed immediately and water molecules layer will be rapidly extruded out and stable binding will be made. Subsequently, barnacle glue secreted from them will immediately tear the surrounding biofilm around the attachment point and establish permanent attachment base, while the corresponding attached biofilm patch of eukaryote genera will be covered by barnacle larvae permanently underneath their bodies.

## 4. Conclusion

The present study was conducted to determine the adhesion efficiency of barnacles on the surfaces of 25 nano-filled coatings. The bacterial communities were determined from the coating biofilm samples for both the protozoan and eukaryotic groups, which were collected from high-adherent and low-adherent coatings. No significant difference was found in the structure of protozoa, while at least two groups in eukaryotes showed significant differences between the two types of coatings. Choreotrichia was proportionally correlated with the adhesion degree of barnacles, while Cryptomonadales was negatively correlated. In addition, two other unidentified eukaryotes had positive correlation with the adhesion degree of barnacles (Figure 3B). These results indicated that eukaryotes had more close relationship with the adhesion process than protozoa did for barnacles. This might be consistent with the observation that the microorganisms mainly composed of eukaryotes retrieved from the attachment plaque (footprint) under barnacle shell. Further studies are needed to explore the molecular and community-level mechanisms that induce the adhesion of barnacle cyprid to specific eukaryotes in the marine biofilm.

## Acknowledgments

This work was funded by the 2018 Weihai Scientific Innovation Project (grant number 2018HW13); the Key research and development plan of Shandong Province (grant number 2016GSF115022) and the Natural Science Foundation of Shandong Province (grant number ZR2018MC002).

